# Is there a trade-off between host generalism and aggressiveness across pathogen populations? A synthesis of the global potato and tomato blight lesion growth data

**DOI:** 10.1101/2022.08.01.502405

**Authors:** Justin SH Wan

**Affiliations:** Institute of Environmental Health and Ecological Security, School of Environment and Safety Engineering, Jiangsu University, Zhenjiang, P.R. China

**Keywords:** Fitness, local adaptation, pathogenicity, *Solanum tuberosum*, *Solanum lycopersicum*, specialisation

## Abstract

*Phytophthora infestans* populations and lineages vary widely in host specificity from specialisation to generalism among potato and tomato. However, generalists that displace others and became dominant on both hosts are relatively uncommon. Generalists may have lower fitness compared to specialists on the host of the latter, which could explain their coexistence at many locations. Lesion size is an aggressiveness metric closely related to fitness in *P. infestans*. A trade-off between generalism and lesion growth rate on the original host can explain the variation among blight populations. I collated the data from cross-inoculation trials on potato and tomato isolates to test for the trade-off. In addition, other metrics related to disease symptoms were included to test whether the degree of specificity is different between populations from potato and tomato, and to explore whether specificity had changed over time. The results indicate a trade-off between generalism and lesion growth rate where higher specificity was associated with significantly faster lesion growth on the original host at the population-level, but not at the lineage-level. Potato and tomato isolates were overall not significantly different in specificity, but tomato isolates tended towards generalism with time. These findings indicate that specialists may avoid displacement on their host by generalists through faster lesion growth, and help explain the common co-occurrence of generalists and specialists. However, a few invasive generalists can rapidly displace competitors and became dominant on both hosts across a broader region. It is likely that important exceptions exist to this trade-off.

## Introduction

Local adaptation to hosts can determine the spatial and temporal population dynamics of plant pathogens (Kröner et al. 2017). Host specificity is indicated by a greater pathogenicity on the original host than on other possible hosts. In the extreme case, the pathogen population is host-specific where the pathogen is only pathogenic on the original host. Adaptive processes occurring within pathogen populations have strong effects on the severity of outbreaks (Sicard et al. 2007). Studying trade-offs and constraints in the population biology of pathogens are of fundamental interest because of their effects on population structure and epidemiology (Burdon, 1993). For example, the emergence of sister species *P. mirabilis, P. ipomaea*, and *P. phaseoli* were associated with specialization onto lesser hosts of *P. infestans* (Raffaele et al. 2010). In addition, severe outbreaks often involve the invasion of new aggressive strains with different properties such as superior performance on hosts or environmental tolerances. Adaptation to hosts is an important aspect of disease emergence (Engering et al. 2013), so it is imperative to study the evolutionary and population biology of emergent pathogens on their hosts.

*Phytophthora infestans* de Bary is a globally distributed heterothallic oomycete pathogen on solanaceous plants. It was the causal agent of the Great Irish Potato Famine that was responsible for millions of deaths. The pathogen was formally described in the 1800s (as *Botrytis infestans* by M.J. Berkeley 1846) and primarily causes disease on potato and tomato on which it is among the most destructive of pathogens (termed late/potato and tomato blight respectively). Populations of newly emergent lineages have occurred continuously, mainly with migration (Shattock 2002; Kildea et al. 2012). Strains vary widely in host specificity among the two host species. The co-occurrence, displacement, and turnover of genotypes through time have been as intriguing to investigators as it was impeding to managers. The global blight populations have undergone several remarkable changes throughout their history, from pan-globally dominant lineages (FAM-1 followed by US-1) to the expansive array of genotypes today (reviewed in Yoshida et al. 2013; Saville et al. 2016). Fitness is related to the effects of pathogens on their hosts and affects the persistence or invasiveness of strains (Pariaud et al. 2009). While no strain persists indefinitely, in invasive strains their persistence is prolonged (e.g., over several seasons or more) and are often dominant (i.e., with high local frequency) across a meta-population level or continental region. Therefore, a better understanding of fitness trade-offs within *P. infestans* on potato and tomato is needed to identify the factors that shape population structure (Andrivon et al. 2013; Saville & Ristaino, 2019).

Host specificity is readily quantified in cross-inoculation trials involving isolates from different host species or varieties. In the literature, aggressiveness is broadly defined as the negative effect that a pathogen has on the host, and can be considered to be any quantitative component of pathogenicity that is non-specific to host genotypes (reviewed in Pariaud *et al*. 2009). In this study, lesion growth rate on host tissue was used as a measure of aggressiveness, which is a life-history trait that is a key predictor of epidemic severity and is widely evaluated across studies on *P. infestans* (Birhman & Singh, 1995; Pariaud et al. 2009; Montarry *et al*. 2010). Previous reports often found that potato isolates (derived from potato as the original host) have a higher preference for potato, but tomato isolates (derived from tomato as the original host) are generalist on both hosts (Legard et al. 1995; Michalska et al. 2016; Kröner et al. 2017). Infectivity on tomato likely evolved relatively recently during initial invasion from the centre of origin in the 1800s, because isolates from the native range were non-pathogenic to tomato (Legard et al. 1995). Blight was long believed to spread from potato to adjacent tomato crops, but not vice versa (Berg, 1926; Small, 1938; Legard et al. 1995; Cohen, 2002). However, more recently it has been revealed that transmission regularly occurs from tomato to potato in relatively new lineages such as 13_A2, US-22, and US-23 (Hu et al. 2012; Frost et al. 2016). Although pathogenicity on tomato likely evolved more recently, the pathogen is typically considered to be as impactful on tomato as it is on potato (Kröner et al. 2017).

Historical observations on host specificity in populations of the pathogen are particularly thought-provoking. *Phytophthora infestans* likely originated from the central highlands of Mexico, where there is a high diversity of tuber-bearing *Solanum* species (Garry et al. 2005; Shakya *et al*. 2018). Prior to the early 1900s blight on tomato was uncommon (Berg, 1926). Reports from the earliest cross-inoculation trials were mostly conducted after 1900, when tomato blight was becoming more prevalent globally. Those reports (between 1900 and 1915) did not mention finding host specificity, or they found that isolates from both hosts were destructive on both hosts under lab and field conditions (Berg, 1926; Oyarzun et al. 1998). However, this changed in the after 1915 when remarkable specificity was frequently noted either on both potato and tomato isolates or in potato isolates only (from Holland, Australia, USA and the UK; Berg, 1926). This was followed by the emergence of US-1 lineage which became dominant from the 1930s. During this period, studies on populations from different regions showed the majority had no remarkable specificity, but potato isolates were still more likely host-specific. For example, Wilson and Gallegly (1955) examined isolates from Scotland, the Netherlands, Canada and the USA, and reported that 8 out of 29 potato isolates were host-specific versus only 2 out of 16 tomato isolates. Isolates from Israel at the time showed similar proportions (9 of 25 potato isolates vs. only 2 of 25 for tomato isolates; Kedar et al. 1959). Other cross-inoculation studies conducted during US-1 dominance in Japan (Kishi, 1962), the USA (before 1970, Flier et al. 2003), and Germany (Günther et al. 1970) also reported little or no specificity. This period was followed by the emergence of an array of successive lineages from the mid- to late-1980s up to the present where US-1 became either extinct or rare relative to other lineages. Studies on surviving (or remaining) US-1 populations after the early 90s overwhelmingly found host specificity in populations from either host species, but mostly on tomato and not potato (Erselius et al. 1997; Oyazun et al. 1998; Vega-Sanchez et al. 2000; Suassuna et al. 2004; Chen et al. 2009). This pattern is reversed In North America where the remaining US-1 potato isolates tended to be host-specific, but not the tomato isolates (Legard et al. 1995; Platt, 1999). Subsequently emerged lineages vary widely in the specificity spectrum. It must be noted that specificity mainly pertains to the population-level rather than the lineage-level (or genotypic-level; Pariaud et al. 2009). For example, in the 90s isolates in Uganda and Kenya from potato and tomato were both US-1, but there were differences in host specificities and glucose-6-phosphate isomerase *(Gpi)* phenotypes among populations on the two hosts (Erselius et al. 1997). In another example, US-8 is a well-known potato specialist but in the late-90s populations were reported on tomato on Prince Edward Island, Canada. Pathogenicity tests on those isolates did not find strong specificity to either host (Daayf and Platt 2003).

The overall evidence suggests *P. infestans* genotypes that are more aggressive and could effectively overcome host defences should be more successful, especially those with a greater lesion size and/or higher sporulation (Gisi et al. 2011). As such those two life-history metrics are closely related to fitness across pathogen genotypes (modelled in Montarry et al. 2010). For instance, the relatively high aggressiveness of US-8 (lesion expansion curve) is likely an important factor in the replacement of US-1 in south-western Canada (Miller et al. 1998). Similarly the more aggressive 2_A1 displaced the less aggressive US-1 in east Africa (Njoroge et al. 2018). In Brazil, the more aggressive BR-1 with much faster lesion growth on potato (but less aggressive on tomato) displaced US-1 on potato only (Suassuna et al. 2004). The 13_A2 lineage was dominant on several continents including Europe and Asia, and had a relatively long presence over many seasons. It is more aggressive than other lineages, and was demonstrated to competitively exclude other lineages in the field, which likely explains its quick expansion across Europe within 3 years (Cooke et al. 2012). Meanwhile, strains that can effectively infect a variety of hosts, including resistant hosts should also be favoured over host-specific strains (Seidl Johnson & Gevens, 2014). Many of the invasive strains that spread quickly to high frequency were aggressive on both hosts, including 13_A2 (Cooke et al. 2012), US-11 (Chen et al. 2009; Daayf and Platt 2003), and US-23 (Saville & Ristaino, 2019). The long-standing question remains: Why aren’t pathogen populations all explicitly both highly aggressive and generalist across hosts?

The relationship between pathogenicity components and fitness is complex (Montarry et al. 2010). Although there is abundant evidence that a faster lesion growth is advantageous for fitness, conversely there have been few suggestions high pathogenicity is costly to transmissibility because hosts are weakened or killed too quickly (Pasco et al. 2016; Mariette et al. 2016). This is in line with the ‘virulence-transmission trade-off’ hypothesis which posits that high host mortality limits transmission (note that the “virulence” term was used to mean quantitative pathogenicity on the host; Acevedo et al. 2019). Models and studies on trade-offs on *P. infestans* often did not consider the effects of alternate host habitats, such as tomato plants or wild *Solanum* species (Montarry et al. 2010; Frost et al. 2016). This is an important aspect since populations on tomato and potato often differ in genotypic composition (Lebreton & Andrivon 1998). A strain that is extremely pathogenic on potato can have slower lesion growth on tomato to compensate an assumed lowered transmission rate on potato if the other host is locally available. Even if high pathogenicity is indeed linked with lower transmissibility, a trade-off between host generalism versus aggressiveness could explain the variation in host specificity among *P. infestans* potato and tomato strains (Thrall & Burdon, 2003; Pariaud et al. 2009). Such a trade-off would be in line with generalists being ‘the jack of all trades but a master of none’ and specialists being ‘the master of some’ (Remold, 2012). It is also evident that few lineages highly aggressive on both hosts exist (i.e., ‘the master of all’ strategy). Perhaps a trade-off between the ability to effectively infect both hosts and aggressiveness on the original host may help explain the scarcity of highly aggressive and generalist genotypes.

A range of cross-inoculation trials on *P. infestans* from potato and tomato have been conducted over the past few decades. Meta-analytic approaches use a standardized within-study effect size that can account for the variation among different metrics used across studies (Chandrasekaran et al. 2016; Acevedo et al. 2019), and have been used effectively in uncovering patterns or processes across studies in plant pathology (Madden & Paul, 2011). It is also a good tool for assessing or comparing the efficacy of different disease control measures across trials (e.g. in pepper blight, Wan & Liew 2020). Here, I collated the global lesion growth rate data to analyse the relationship between host specificity and aggressiveness. In addition I tested for differences in specificity and lesion growth rate among potato and tomato isolates. The following predictions were tested: (1) There will be a positive relationship between host specificity and lesion growth rate on the original host; (2) overall, potato isolates will have higher specificity for potato than tomato isolates for tomato; (3) host specificity will tend to reduce over time (with selection for generalists); and (4) isolates of the old US-1 lineage will have slower lesion growth rate than those of successive, more aggressive lineages.

## Methods

### Data collation

To identify studies performing controlled cross-inoculations using potato and tomato isolates of *P. infestans*, I searched ISI Web of Science (Clarivate) and Google Scholar (https://scholar.google.com) in January 2022 using the following terms: ‘Phytophthora infestans’ AND ‘tomato’ OR ‘potato’. Only trials conducted on potato and tomato hosts were selected. Studies that tested pathogenicity on only one out of these two hosts were excluded, as well as studies that did not report the original host of isolates (or was indeterminable based on available information). As such, ‘potato isolates’ or ‘tomato isolates’ refer to the respective original host. To address predictions (2), (3) and (4), studies reporting pathogenicity metrics other than lesion growth rate were included to assess the degree of specificity across populations. Studies must describe some quantitative or semi-quantitative measure of pathogenicity (e.g. lesion size, AUDPC, degree of infection). Results not in journal articles such as research thesis and reports were accepted to reduce publication bias (Madden & Paul 2011). Thus, those reporting only qualitative pathogenicity (i.e. pathogenic or not pathogenic) are excluded. Studies that tested multiple isolates were pooled to one data point (separately for each original host). This was done to avoid pseudoreplication leading to bias towards studies reporting more detailed data.

From each study, I recorded: (1) the original host (potato or tomato); (2) lesion growth rate (hereafter ‘LGR’ in millimetres per day) on both hosts. Alternatively another measurement of pathogenicity was recorded to address the other predictions (2-4). Semi-quantitative metrics such as percentages, disease severity scores or scaled grading were also accepted as host specificity was analysed using non-parametric analyses. For studies reporting lesion areas, the length should ideally be assumed elliptical and calculated using width (length = area / [0.25π · width]; Vleeshouwers et al. 2000). However, the width of lesions was not reported in the studies, so lesions were assumed to be square-shaped and converted using: length = √mm^2^. divided by the number of days post-inoculation to obtain mm/day; (3) location of the pathogen population; and (4) time of isolation from hosts (calendar year). The mean calendar year was used where multiple isolates were pooled.

The within-study differences in pathogenicity of isolates on the two hosts were standardized using log10 response ratio (LRR) as the effect size. Very negative LRR values (less than minus one) are associated with extreme specificity (i.e., non-pathogenic on the other host). Conversely, a very positive LRR (greater than one) indicates the pathogen is only pathogenic on the other host and therefore the original host is likely a sink habitat. This standardized LGR metric was used to gauge the aggressiveness of pathogen populations on the two hosts. Henceforth, *LGR_origin_* and *LGR_other_* refer to the LGR on the original host and the other host respectively. *LRR_LGR_* refers to the LRR calculated from LGR, used in the analyses on aggressiveness.

To test the relationship between specificity and aggressiveness, the analysis on *LGR_origin_* vs. *LRR_LGR_* is divided into two parts: 1) Population-level patterns. By extracting only one data point per host from each study (i.e., pooling isolates and lineages from each host), each data point is representative of the whole population on the hosts at that location. Thus, in this part I test whether host specificity is associated with greater aggressiveness across pathogen populations strictly in general. 2) Lineage-level patterns (sub-population). Studies reporting the LGR of multilocus genotypes (MLG) or lineages were recorded as separate entries in another dataset. This allows for analysis on whether MLGs with higher specificity also tend to be more aggressive. Where only a single lineage is found on a host the pathogen population is represented by that lineage on that host.

In addition, I compared the host specificity and LGR of US-1 isolates and other lineages. From all the studies collected from the literature search, studies that tested US-1 isolates were identified. Part of these studies also tested isolates that are not US-1 along with other isolates (which were pooled with US-1 in the other analyses). Those were un-pooled and entered as separate entries comprising of ‘US-1’ and ‘Other’ lineage categories. Limiting the analysis to studies that tested both US-1 and successive lineages within the same trial allows for more direct comparison.

### Statistical analyses

Normality in distributions was checked using quantile-quantile plots (Q-Q plots) prior to analysis, which indicated the LRR data did not follow a normal distribution. To assess whether host specificity is associated with greater LGR, the relationship between *LGR_origin_* and *LRR_LGR_* was analysed in two ways. For the population-level analysis, a median-based linear model with the Thiel-Sen single median method was used (Komsta & Komsta, 2013). Theil-Sen estimates the slope from the median among all possible combinations of slopes between points. This method is robust and assumes all LGR values are different, so all duplicate values was adjusted by a token amount (0.001). For the lineage-level analysis, the median model could not account for non-independence from data points extracted from the same study (there were only 10 studies reporting the lineage and data points from the same study ranged from 2-7), so linear mixed model was used with ‘Study’ as a random effect. The *LRR_LGR_* data was exponential transformed using exp(*LRR_LGR_*). Three outliers with *LRR_LGR_* < - 1 were removed prior to analysis.

For the analysis on whether potato and tomato isolates had statistically significant host specificities, a non-parametric bootstrap approach was used to estimate the mean effect size (LRR) and confidence intervals (bootstrapped 95% CI). Bootstrapping is based on the method used in van den Noortgate and Onghena (2005) and involves random subsampling with replacement over 1000 iterations. This approach is more suitable because pathogenicity trials often did not report the variance (as compared to a classical meta-analysis approach). Non-parametric bootstrapping does not emphasize thresholds for statistical significance, but emphasizes effect size and confidence intervals (Rillig et al. 2019). Median-based linear models were used to test the relationships between *LGR_origin_* vs. *LRR_LGR_*, and LRR vs. time (calendar year). The LRR and LGR differences between potato versus tomato isolates, and US-1 versus other lineages were analysed using Mann-Whitney U tests.

All analyses were conducted using R (v.3.6.1, R Core Development Team 2019). Bootstrapping was conducted using the *boot* package (v. 1.3-22) and the *metaphor* package (v. 2.1-0) was used to calculate effect sizes. The *mblm* package (v.0.12.1; Komsta & Komsta, 2013) was used to run the median-based models, and the *lmer* package was used to run the linear mixed model on the lineage data. Host specificities (LRR) were considered statistically significant where the bootstrapped CI do not overlap with zero (e.g., Chandrasekaran et al. 2016). GetData Graph Digitzier software (v. 2.26) was used to extract data from figures (available at getdata-graph-digitizer.com).

## Results

This search included 78 host specificity comparisons of *P. infestans* from 44 studies in total (*k* = 44 and 34 for potato and tomato respectively, Fig. 1). Of these, 43 were of lesion size data. Most studies (32 of the 44 studies) conducted cross-inoculation trials on isolates from both hosts within the same trial. The data was drawn from all over the globe and all inoculation trials were conducted under controlled trial conditions (Fig. 2). Ten of the studies tested US-1 isolates along with isolates of successively emergent lineages. An outlying data point with LRR value of greater than 1 (i.e., non-pathogenic on the original host but highly pathogenic on the other host) was removed from all analyses (summary statistics in Fig. S1). For aggressiveness (LGR), all data were derived from detached leaf assays on potato and tomato. For the expanded host specificity data used in comparing specificity (LRR) within and among potato and tomato isolates, the vast majority of pathogenicity metrics were directly related to lesions (63 out of 78 data points, e.g., lesion size, growth, percentage, or number), and 5 data points were based on the evaluation of lesions (proportion of plants or leaves with lesions or disease ratings /grading based on the type of lesions – sporulating, expanding or non-expanding). The remainder (10 data points) include sporulation (density or proportion of lesions with sporulation – 5 data points) and overall disease severity (5 data points). The inoculation approach used was mostly detached leaf assays (64 out of 78 data points, with one combining data from detached stem and leaf inoculations), 4 data points were of leaf inoculation in planta, 10 used spray inoculation and a single one was of tuber and stem inoculation. The isolation time of pathogen isolates from plants in the field ranged from 1970 to 2017. The full dataset and references are provided in the Supplementary Material (Table S1).

**Fig. 1.** Flow diagram of the literature search and the screening process, detailing the number of studies excluded during screening up to the final number of studies included.

**Fig. 2.** World map detailing the locations of *Phytophthora infestans* potato and tomato populations tested in cross-inoculation trials. Symbols with lighter shades represent imprecise locations specified to the general region only. Geographically distant populations tested within the same study are joined by dotted lines.

At the population-level, there was a significant positive relationship between host specificity (*LRR_LGR_*) and aggressiveness on the original host, *LGR_origin_* (*P* < 0.05, Table 1a, Fig. 3), but not at the lineage-level (Table 1b). These together indicate that host specificity is generally associated with faster lesion growth across pathogen populations across locations and regions, but not among genotypes. Extreme specificity to potato or tomato was rare, and pertains to only few cases (such as certain US-8 populations; Table S1). Overall, potato and tomato isolates had statistically significant specificities for their original host (LRR estimates with 95% bootstrap CI: −0.26 [−0.09, −0.52] and −0.18 [−0.01, −0.41] respectively, *P* < 0.05; Fig. 4). This means that overall the pathogenicity when infecting the other host is reduced to ~55% and 66% of when on the original host for potato and tomato isolates, respectively. The specificity difference between potato versus tomato isolates (LRR −0.26 vs. −0.18 respectively) was not significant (W = 673, *P* = 0.45; Fig. 4a), and neither were the differences when only the LRR based on lesion growth was considered, *LRR_LGR_* (W = 173, *P* = 0.18). There was also no difference in aggressiveness (LGR) between potato and tomato isolates on their respective original hosts, *LGR_origin_* (W = 239, *P* = 0.99; Fig. 4b, c). There was no significant relationship between LRR and isolation time for potato isolates (*P* = 0.54), but for tomato isolates there was a significant positive relationship, where the median LRR increased from −0.18 (i.e. reduced to ~84% pathogenicity on the other host compared to when on original host) to 0.06 (i.e. to ~106% pathogenicity on other host compared to when on original host) over around 30 years (*P* < 0.01, Table 2; Fig. 5). This suggests that the most recent tomato populations have little if any specificity – from year 2010 onwards the tomato isolates data is mainly represented by generalists such as US-22, US-23, and 13_A2.

**Fig. 3.**
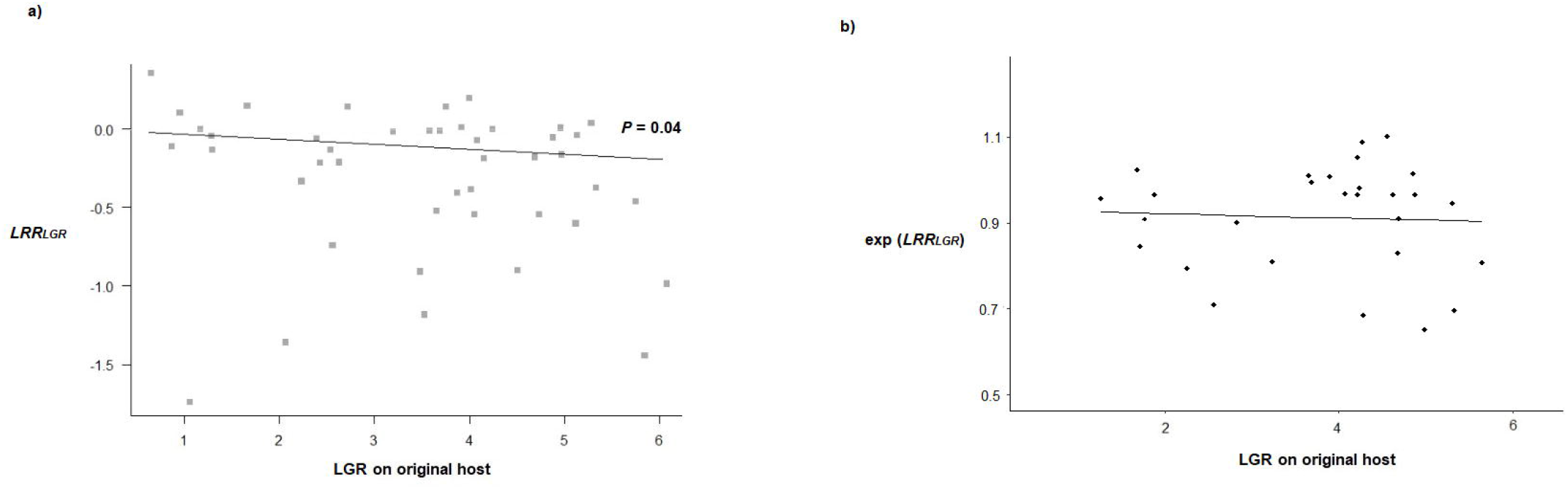
Relationships between host specificity based on lesion growth rate (*LRR_LGR_*) and lesion growth rate on the original host (*LGR_origin_*) for potato and tomato isolates of *Phytophthora infestans* at the: a) Population-level, where each data point is representative of the pathogen population present on each host at a location. Populations with no specificity are indicated by *LRR_LGR_* = 0; b) Lineage-level, with each data point of the exponential transformed *LRR_LGR_* representing a lineage tested within a given study. Lineages with no specificity are indicated by exp(*LRR_LGR_*) = 1. More negative LRR values are associated with greater specificity on the original host in all cases.

**Fig. 4.**
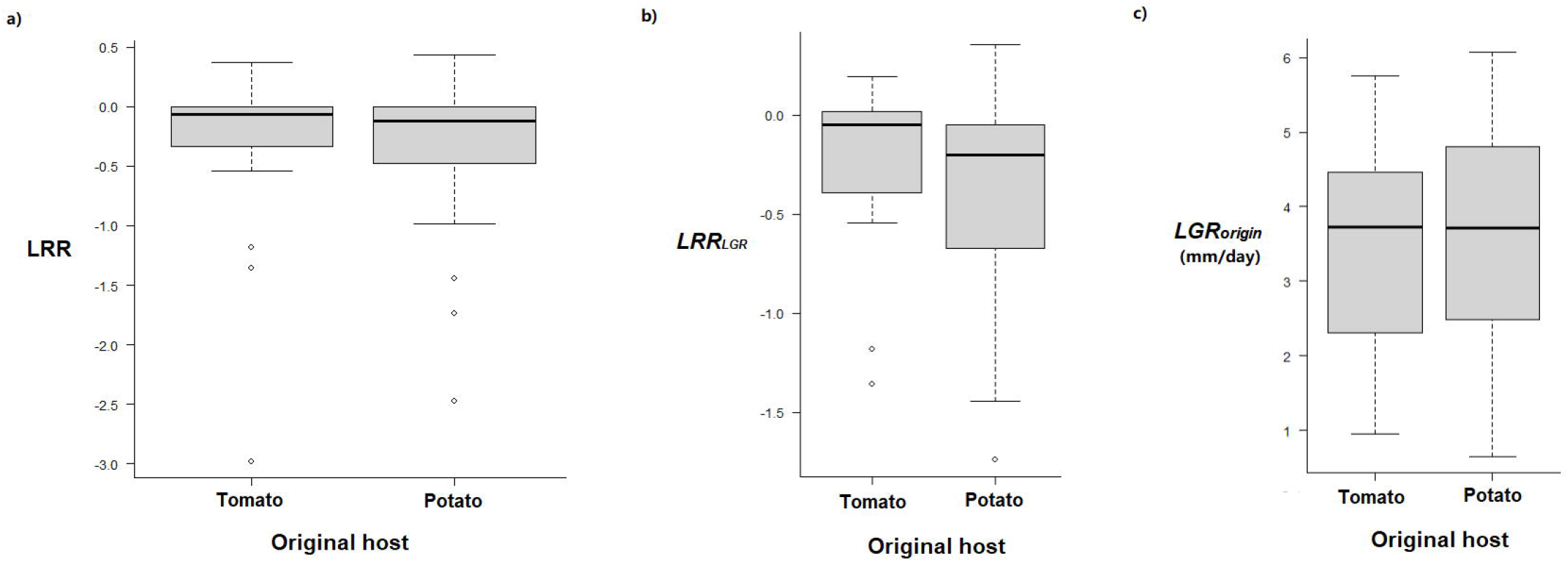
Boxplots comparing *Phytophthora infestans* populations from potato and tomato, showing the 0.25, median (0.5) and 0.75 quartiles: a) The overall degree of host specificity (LRR) as indicated by different pathogenicity metrics; b) host specificity in the subset as calculated from lesion growth rate (*LRR_LGR_*), and c) lesion growth rate on the original host (*LGR_origin_*).

**Fig. 5.** Relationship between host preference (LRR), as indicated by different pathogenicity metrics, versus isolation time (calendar year) in *Phytophthora infestans* a) potato and b) tomato isolates, where each data point represents a pathogen population present during the time period.

**Table 1.** Results on aggressiveness on the original host (*LGR_origin_* in millimetres per day) versus host specificity (*LRR_LGR_*) at the: a) Population-level from median-based model (*k* = 43), and b) lineage-level from mixed effects linear model with ‘Study’ as a random effect level. *LRR_LGR_* was exponential transformed for normality in the residuals (*k* = 28). Three outliers *LRR_LGR_* < −1 were excluded). Bold values indicate statistically significant effects (*P* < 0.05).

| Source | Estimate | Median absolute deviation | V | P |
| --- | --- | --- | --- | --- |
| Intercept | -0.00 | 0.21 | 470 | 0.78 |
| $LRR_{LGR}$ | -0.03 | 0.10 | 316 | **0.037** |

b)
| Source | Estimate | df | t | P |
| --- | --- | --- | --- | --- |
| Intercept | 0.93 | 26 | 12.41 | **<0.001** |
| $\exp(LRR_{LGR})$ | -0.03 | 26 | -0.26 | 0.797 |

**Table 2.** Results from median-based linear models on host specificity (LRR) versus isolation time (calendar year) in *Phytophthora infestans* potato and tomato isolates. Bold values indicate statistically significant effects (*P* < 0.05).

| Source | Estimate | Median absolute deviation | V | P |
| --- | --- | --- | --- | --- |
| Potato isolates ($k = 40$) | | | | |
| Intercept | -0.41 | 10.29 | 448 | 0.62 |
| Calendar year | 0.00 | 0.01 | 328 | 0.54 |
| Tomato isolates ($k = 32$) | | | | |
| Intercept | -17.00 | 26.98 | 126 | **0.009** |
| Calendar year | 0.01 | 0.01 | 402 | **0.009** |

Focussing on US-1 versus successively emerged lineages from the studies, the sample size available for analysis was relatively low (*k* = 11 and *k* = 10 for US-1 and other isolates respectively) and fewer reported lesion growth rates (*k* = 7 and *k* = 6 respectively). Two US-1 tomato isolates had extreme specificity for potato and was non-pathogenic on its original host which were removed from the analyses (LRR 2.31 and 1.58). There were no significant differences in LRR among isolates of US-1 and other lineages (LRR −0.22 vs. −0.30 for US-1 and other lineages respectively; W = 51, *P* = 0.66). Furthermore, there was no significant difference in *LGR_origin_* among US-1 and other successively emergent lineages (LGR 4.12 vs. 3.47 mm per day respectively; W = 5, *P* = 0.57). The low availability of data between US-1 and new isolates means that the results for this comparison should be taken with caution. Nonetheless as the analyses are on a subset that assessed US-1 and other lineages within the same controlled study, it is indicated that those US-1 isolates do not necessarily have lower LGR than those of newer lineages.

## Discussion

The current synthesis of the global data suggests generalism tended to be costly to lesion growth rate in *P. infestans* across pathogen populations on each host. *Phytophthora infestans* is a hemibiotrophic pathogen with a clear necrotrophic phase, so aggressiveness should be under selection in the absence of trade-offs (Jarosz & Davelos, 1995; Montarry et al. 2007). Aggressive lineages often succeed less aggressive ones (e.g. Miller et al. 1998; Cooke et al. 2012), so a slower lesion growth rate is likely a fitness cost for the ability to effectively infect both hosts for many generalist populations. From the data, the most aggressive populations (with LGR on original host > 5 mm/day) were mostly host-specific (5 out of 7 data points had *LRR_LGR_* < −0.30). Moreover the top two (with LGR on original host ~6 mm/day were extreme in host specificity (*LRR_LGR_* < −1, see Fig. 3). However, this trade-off was not found at the lineage-level. Populations typically consist of multiple competing lineages. Within each population, specialists may only require a small fitness advantage on its host (e.g. a slightly higher LGR) relative to co-occurring generalists to avoid displacement by them (Kröner et al. 2017). This difference may be too small to be detected in the lineage data, or that there is a lot of variance in the data. Unfortunately, there were only 10 studies that reported lineage specific data, and other studies combined data from different lineages or did not identify the MLG of isolates, which limits the certainty in the conclusion. This main finding indicates that populations consisting of host-specific genotype(s) tend to be more aggressive on that host versus populations that comprise of more generalists. This trade-off is in line with the classic trade-off between virulence and aggressiveness in the GFG model within a given host species (Thrall & Burdon, 2003), and may help explain the existence of specialists when both hosts are often abundantly available in the environment. The finding is consistent with the first prediction that there will be a positive relationship between host specificity and LGR on the original host.

There are some genotypes where this trade-off may not be applicable. Invasive generalists can quickly displace competitors and became dominant on both hosts across a greater region, such as 13_A2 across Asia (Dey et al. 2018). In those cases the whole populations on both hosts is represented by the single lineage. Such dominant lineages are relatively few because the vast majority of lineages are not and do not become dominant. There are a number of possibilities for this. Migration restrictions and control measures (e.g. biosecurity on potato trade) can prevent introduction to new areas and inhibit their spread. Other explanations include limitations in genetic mechanisms leading to the emergence of aggressive generalists (e.g. epistatic pleiotropy, Remold, 2012) or compensatory selection that may recover the costs of generalism (Pariaud et al. 2009). The genetic mechanisms and ecological context for how highly aggressive generalists arise remain to be explored (Burdon, 1993). However whether those invasive genotypes are more aggressive than most specialists could not be interpreted from the data. While the current study found that populations with faster lesion growth tended to have higher specificity through the standardised LGR, it is not suitable to directly compare specific data points sourced from different studies for inferences on their relative pathogenicity or fitness. This is because of variation among conditions connected to each individual trial (e.g., host cultivar, culturing conditions, environmental parameters etc.) that replication may produce different results (Andrivon et al. 2013). For instance, the widespread 13_A2 from India that was dominant on both hosts only had a moderate lesion expansion rate in trials (~3.5 mm/day in Chowdappa et al. 2015; ~1.6 to 2.6 mm/day in Dey et al. 2018). It is possible that a specialist more aggressive (on its host) than an invasive generalist could persist on that host. Testing isolates together under controlled conditions (e.g. with competitive interactions) is still necessary to measure the relative aggressiveness of genotypes.

Several factors could potentially explain why generalist populations tended to have slower lesion growth than specialists on the same host. First, specialisation may be associated with increased fitness in that environment (e.g., more effective suppression of host defences), regardless of any fitness costs associated with generalism (i.e. ‘the jack of all trades’ strategy; Remold, 2012). Second, genetic change involved in adaptation to alternative hosts may involve fitness costs due to antagonistic pleiotropy. This includes costs associated with virulence genes to overcome host resistance factors (Pariaud et al. 2009; Montarry et al. 2007, 2010). Third, specialists should have a lower accumulation of deleterious alleles (Whitlock, 1996), which may hinder the competitiveness of a generalist in the presence of specialists (Kawecki, 1994). Yang et al. (2016) found no evidence that mutation accumulation was involved in a trade-off between thermal tolerance and pathogen growth rate in *P. infestans*, but may be driven by selection under antagonistic pleiotropy. Virulence was traditionally defined as the ability of a pathogen to overcome host defences, especially in the gene-for-gene (GFG) model so that high virulence is associated with the ability to infect more hosts (Laine & Barrès, 2013).

Even though lesion size is regularly used to gauge aggressiveness and the severity of outbreaks (e.g. Miller et al. 1998; Pariaud et al. 2009), both LGR and sporulation capacity are closely linked to fitness (Montarry et al. 2010). Thus considering LGR alone may not accurately reflect aggressiveness in cases where sporulation capacity (i.e. spore density within lesion area) is the key factor. There was insufficient data for analyses based on sporulation. In the compiled literature, there were only 29 data points in total for sporulation, and measurements differed in scaling (12 were scaled to area or length and 17 were scaled to volume, Table S1). Other metrics (e.g. host mortality and infection efficiency) were even less commonly reported. Some lineages had fast lesion growth on both hosts but low sporulation density on one host (i.e., the host it is less often isolated from). An example where sporulation is more important is in 23_A1 from Algeria that is mainly found on tomato (Beninal et al. 2022). A trial found that it has relatively fast lesion growth on both hosts (4.9 – 5.0 mm/day) but lower sporulation capacity on potato (Belkhiter et al. 2019). This is especially compared to its specialist competitor (13_A2 on potato), which had a slightly lower LGR (4.7 mm/day) but superior sporulation on potato (see Figure 2a of Belkhiter et al. 2019). Conversely in North America (around 2010), US-23 is genetically similar to 23_A1 but caused major epidemics on both hosts (Danies et al. 2013). It has both higher sporulation capacity and LGR than competing lineages US-22 and potato specific US-24, except US-8 on potato (Danies et al. 2013). A few years later it completely displaced US-22 and US-24 (Saville & Ristaino, 2019). Whether sporulation capacity is involved in trade-offs could be addressed in future research. Nonetheless genotypes with high LGR on both hosts should be considered potentially invasive.

Interestingly, genotypes with weaker pathogenicity may sometimes persist. There was a suggestion that potato specialist US-24 may persist in more northern locations despite being less aggressive than generalists US-22 and US-23 due to its superior performance under cold conditions (Danies et al. 2013). The genetically similar potato-specific US-8 was almost entirely displaced but some remaining isolated populations (mainly in the West Coast of the USA) may have been temporarily sustained by inoculum sources at certain localities (Saville & Ristaino, 2019). Long-term studies taking account of the complete life cycle of the pathogen conducted over multiple seasons are likely needed to more reliably assess fitness (Andrivon et al. 2013). Other considerations that may influence the current results include the effects of fungicide presence and other environmental variation (e.g. temperature) that were not replicated in lab trials. There are alternative metrics such as infection efficiency and competitive ability (Lebreton et al. 1999) that could not be addressed by this study that may also be important in gauging aggressiveness.

Contrary to the second hypothesis that potato isolates will be more host-specific, potato and tomato populations did not differ significantly in specificity, but there may be around twice as many highly host-specific populations for potato than for tomato by proportion (12 out of 44 data points or 27% for potato, versus 4 out of 34 data points or 12% for tomato) as defined by an LRR of less than −0.5 (i.e. pathogenicity reduced to < 32% compared to infecting original host; Fig. 5). The reduction in host specificity of pathogen populations on tomato over the last few decades is interesting, and only partially supports the third hypothesis (that overall host specificity will reduce with time) since the specificity of potato isolates did not change over time. Together, these results generally support the long-held belief that tomato isolates are more generalist than potato isolates. This may be particularly relevant for cold regions where potato isolates could survive on tubers but tomato isolates may need other hosts, so specialisation to tomato may not be under selection.

In the current study, lower lesion growth rate on the original host or higher specificity in US-1 populations was not found. Past studies often observed the displacement of US-1 by more aggressive lineages (Legard et al. 1995; Reis et al. 2003; Suassuna et al. 2004; Chen et al. 2009). The non-significance difference in lesion growth could be due to the low number of data points available in this case. Alternatively there is a possibility that the persistent US-1 reflects selection on a subset equally aggressive as newer invading populations. This survivorship effect may have also been observed in *P. cinnamomi*, where the A2 mating type is predominant and displaced the less virulent endemic A1 throughout Asia (Arentz, 2017). The remaining extant *P. cinnamomi* A1 has similar aggressiveness (i.e. lesion size) to the A2 populations (Dudzinski et al. 1993; Robin & Desprez-Loustau, 1998). Nonetheless, future routinized controlled trials testing the pathogenicity (especially of LGR and sporulation capacity) of pathogen populations on the two hosts would be greatly beneficial for the management of extant and emergent lineages of potato and tomato blight (such as 36_A2 and 41_A2 in Europe; EuroBlight network: <u>agro.au.dk/forskning/internationale-platforme/euroblight</u>, accessed Nov 2022), particularly under the effects of climate change (Pangga et al. 2011)..

## Conclusions

The current synthesis revealed a trade-off between generalism and aggressiveness across *Phytophthora infestans* populations on potato and tomato. Overall, populations consisting of more specialists have faster lesion growth rate on their original host than populations consisting of generalists on the same host, which may help explain the patterns in population composition of generalists and specialists among the two hosts. However, this trade-off may not apply for few aggressive and invasive generalists that became dominant on both hosts across a broader region. Although the level of specificity among potato and tomato isolates was not significantly different, there was an indication tomato isolates tended to become more generalist over the last few decades. Due to the variability in how pathogenicity traits relate to fitness, long-term studies (e.g. over multiple seasons) are likely needed to better understand the trade-offs involving pathogenicity. The trade-off between generalism and aggressiveness may help explain the regular coexistence of specialist and generalist lineages at many locations.

## Statements and Declarations

The sole author declares there is no conflict of interest.

## Data availability statement

All data generated or analysed during this study are included in this published article (and its supplementary information files).

## Acknowledgements

The author expresses gratitude to the researchers whose data contributed to this synthesis. Special thanks go to Susan Rutherford for her support and help proofreading the manuscript and Edward Liew for the earlier discussions on Phytophthora and for his advice. Justin SH Wan was supported by the Jiangsu University Science Foundation Fund (20JDG056).

## Supplementary materials

**Table S1**

The datasets used in the synthesis of host specificity and lesion growth rate in *Phytophthora infestans* pathogen populations and lineages on potato and tomato.

**Fig. S1**

Summary median and quantiles for host specificity (LRR) among potato and tomato populations of *Phytophthora infestans* in global regions (*k* = sample size). More negative LRR values indicate higher specificity. Very negative values (such as less than −1) are host-specific to the original host. One population had a very positive LRR and was host-specific to the other host which was excluded from all analyses (indicated by the grey arrow). A value of zero indicates identical pathogenicity on both hosts indicating no specificity.

## Notes

### Competing Interest Statement

The authors have declared no competing interest.

### Summary of Updates

New additional analyses.

