## Supplementary figures and images for "Is there a trade-off between host generalism and aggressiveness across pathogen populations? A synthesis of the global potato and tomato blight lesion growth data"

### Fig. S1

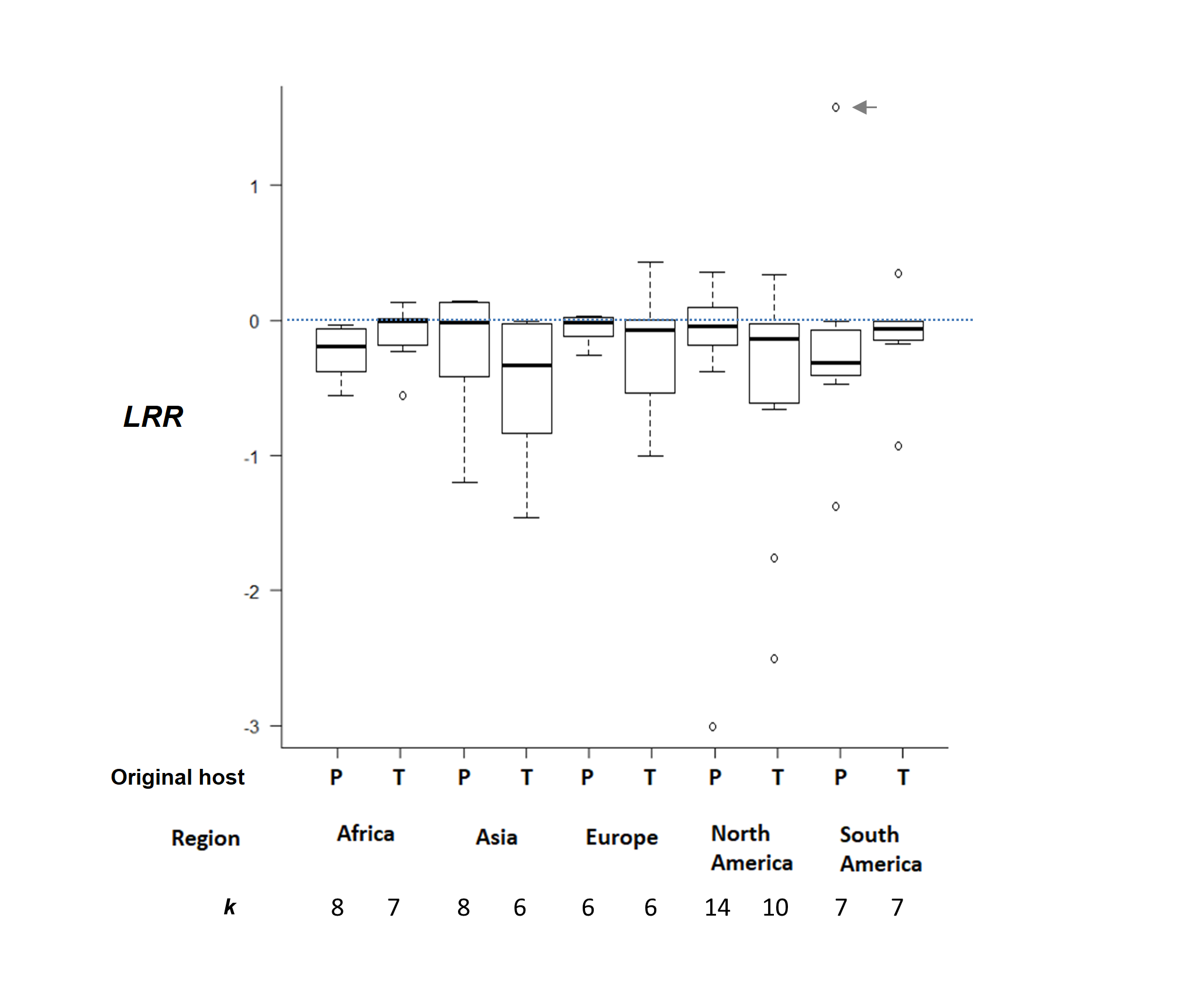
